# Proteomic Features of Adolescents and Young Adults with Soft Tissue Tumours

**DOI:** 10.1101/2023.11.18.567667

**Authors:** Yuen Bun Tam, Kaan Low, PS Hari, Madhumeeta Chadha, Jessica Burns, Christopher P Wilding, Amani Arthur, Tom W Chen, Khin Thway, Anguraj Sadanandam, Robin L Jones, Paul H Huang

## Abstract

Adolescents and young adult (AYA) patients with soft tissue tumours (STT) including sarcomas are an underserved group with disparities in treatment outcomes. To define the molecular features between AYA and older adult (OA) patients, we analysed the proteomic profiles of a large cohort of STT across 10 histological subtypes (AYA n=66, OA n=243). AYA tumours are enriched in proteins involved in mitochondrial metabolism while OA patients have elevated inflammatory and cell cycle signalling. By integrating the patient-level proteomic data with functional genomic profiles from sarcoma cell lines, we show that the mRNA splicing pathway is an intrinsic vulnerability in cell lines from OA patients and that components of the spliceosome complex are independent prognostic factors for metastasis free survival in AYA patients. Our study highlights the importance of performing age-specific molecular profiling studies to identify risk stratification tools and targeted agents tailored for the clinical management of AYA patients.

## Introduction

Soft tissue tumours (STT) are rare mesenchymal tumours that span >80 histological subtypes of distinct biology and genetics^1^. These include malignant cancers such as sarcomas as well as benign diseases such as desmoid tumours (DES). Sarcomas have a higher incidence in the adolescent and young adult (AYA) age group (15-39 years at the time of diagnosis, 8% of all cancer diagnosis), compared to older adults (OA) (>39 years, 1% of all cancer diagnosis)^2,3^. Despite the increased incidence, improvements in the survival in AYA patients with soft tissue sarcomas (STS) have lagged behind other age groups^4^. Reasons for this disparity are multi-factorial, and include under-representation in clinical trials^4,5^, unique psychosocial considerations^6,7^, inadequate age-specific services^8-10^, and poor knowledge of their unique biology^11,12^. A recent study showed that STS histological subtypes typically sensitive to chemotherapy in other age groups are instead chemoresistant in AYA patients^13^, which suggests that patients in this age group may have distinct biological differences compared to either OA or paediatric patients. In the case of non-rhabdomyosarcoma STS (NRSTS) where the majority of current treatment guidelines relies on drugs that have been optimised in OA patients^14,15^, the lack of therapies tailored to AYA patients is a major unmet need and a key barrier to improving survival rates.

Several recent pan-cancer analyses have leveraged on publicly available datasets (TCGA, ICGC, GENIE) to demonstrate that there are age-associated genomic, transcriptomic and immune microenvironmental differences across multiple cancer types that include STT^16-20^. For instance, Lee et al., showed that sarcoma patients <50 years old had lower immune-related pathway expression compared to patients >50 years of age at the transcriptomic level^17^. They further determined that at the genomic level, the older sarcoma patients had higher copy number variation rates. However, the AYA age group is under-represented in all these studies with only a small number of STT patients and histological subtypes included. Furthermore, these aggregate analyses do not consider well-established differences in the spectrum of STT histological subtypes in AYA versus OA patients^21^. Prior studies that have undertaken molecular profiling of AYA sarcoma specimens including a recent EORTC SPECTA-AYA study have focused exclusively on genomic and transcriptomic data^20,22,23^. While informative, these technologies do not provide a direct measure of proteins which are key mediators of tumour cell signalling and the largest class of targets for oncology drugs^24-27^, making it challenging to bridge the translational gap towards clinical applications. Given the unique tumour, microenvironmental and host differences between AYA and OA patients^12^, it is likely that AYA patients with STT harbour distinct molecular features which may influence clinical and treatment outcomes, although this has yet to be conclusively demonstrated. Due to the rarity and heterogeneity of STT, to date there are few studies that have systematically evaluated the molecular differences between AYA and OA patients.

Here we undertake a detailed analysis of the proteomic features in AYA and OA patients across 10 histological subtypes of STT. By interrogating clinically annotated proteomic profiles in a large cohort of AYA and OA patients and undertaking integration with functional genomics data derived from sarcoma cell lines within the Cancer Cell Line Encyclopaedia (CCLE), we demonstrate that there are significant differences in the biological networks and intrinsic vulnerabilities between these two age groups with implications for biomarker development and therapy selection.

## Results

### Cohort and clinicopathological data

The cohort comprises primary tumour specimens from 309 patients (AYA=66, OA=243) for which comprehensive proteomic profiles by mass spectrometry (MS) have previously been generated by our laboratory^28^. Nine sarcoma subtypes are represented, including alveolar soft part sarcoma (ASPS), angiosarcoma (AS), clear cell sarcoma (CCS), dedifferentiated liposarcoma (DDLPS), desmoplastic small round cell tumour (DSRCT), epithelioid sarcoma (EPS), leiomyosarcoma (LMS), synovial sarcoma (SS) and undifferentiated pleomorphic sarcoma (UPS) (full clinicopathological information provided in Table S1). Additionally DES, a benign locally infiltrative STT with no metastatic potential and a relatively high incidence in the AYA age group, was included in the cohort^29^. When broken down by age groups, AYA patients are enriched for ASPS (100%) and DSRCT (75%), while OA patients are enriched for UPS (98%), DDLPS (95%), LMS (91%), AS (90%) and CCS (67%) (Figure 1A). There are almost equal number of AYA and OA patients in SS (44% AYA), DES (49% AYA) and EPS (50% AYA). The cohort has a female predominance (male [37%], female [63%]), with a broad distribution of different anatomical sites in each age group (Figure 1A). Consistent with a previous study of a large cohort of ∼5000 STS patients by the Scandinavian Sarcoma Group (SSG)^3^, our cohort has a higher proportion of patients with grade 3 (OA: 53%, AYA: 15%) and large tumours ≥15cm (OA: 26%, AYA: 12%) in the OA age group (Table S1). Similar to the SSG study, univariate Cox regression analysis showed that patients belonging to the AYA age group were at a statistically significant lower risk of death compared to patients in the OA group (Hazard Ratio (HR) =0.423, 95% Confidence Interval (CI) = 0.228-0.786, p = 0.0065) (Figure 1B). There was no statistically significant difference in the two age groups for metastasis free survival (MFS) and local relapse free survival (LRFS) (Figure S1). Note that DES was not included in any the survival analysis undertaken in this study because they are not malignant.

**Figure 1.**
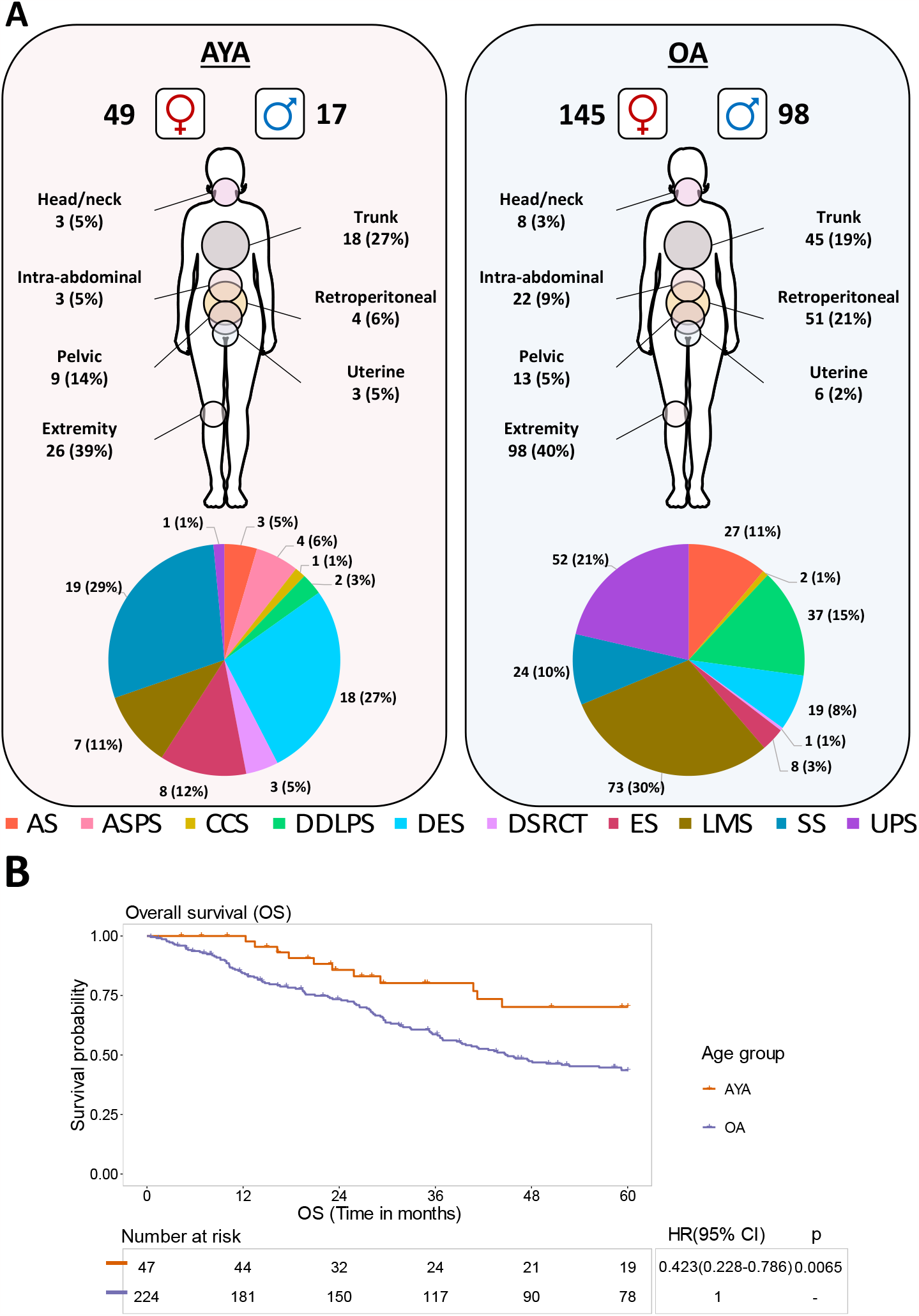
Overview of the AYA and OA patients in the cohort. (a) Distribution of patient sex, anatomical site, and histological subtype within the adolescent and young adult (AYA) and older adult (OA) cohorts. (b) Kaplan–Meier plot of overall survival (OS) for AYA and OA patients. Hazard ratio (HR), 95% confidence intervals (CI) and p-value determined by univariable Cox regression. AS = angiosarcoma, ASPS = alveolar soft part sarcoma, CCS = clear cell sarcoma, DDLPS = dedifferentiated liposarcoma, DSRCT = desmoplastic small round cell tumour, DES = desmoid tumour, EPS = epithelioid sarcoma, LMS = leiomyosarcoma, SS = synovial sarcoma, UPS = undifferentiated pleomorphic sarcoma.

### Analysis of the AYA and OA proteomic landscape

A total of 8148 proteins were identified with 3299 proteins quantified across all samples (Figure 2A and Table S2). We defined the proteins that are significantly upregulated in AYA versus OA patients. Following multiple testing correction, 32 and 35 proteins were identified to be significantly upregulated in AYA or OA respectively (FDR < 0.05, fold change > 2) (Figure 2B and Table S3). Our analysis finds that OA patients harboured a significant upregulation of proteins involved in DNA replication (MCM complex), cell cycle regulation (CDK1, CDKN2A) and immune-regulation (CD163, B2M, IL4I1) while AYA patients displayed an upregulation of proteins involved in mitochondrial metabolism (NDUFA9, SUCLA2, FDXR and ACADVL) and skeletal and cardiac myosin chains (MYL1, MYL2, MYLPF and MYH7). Multivariable logistic regression analysis was performed to adjust for potential confounding factors (tumour size, grade, anatomical site, performance status and histological subtype) which led to five proteins remaining significant between the two age groups (AYA: NDUFA9, SUCLA2, TUBB2B, MACROH2A2 and OA: CDK1).

**Figure 2.**
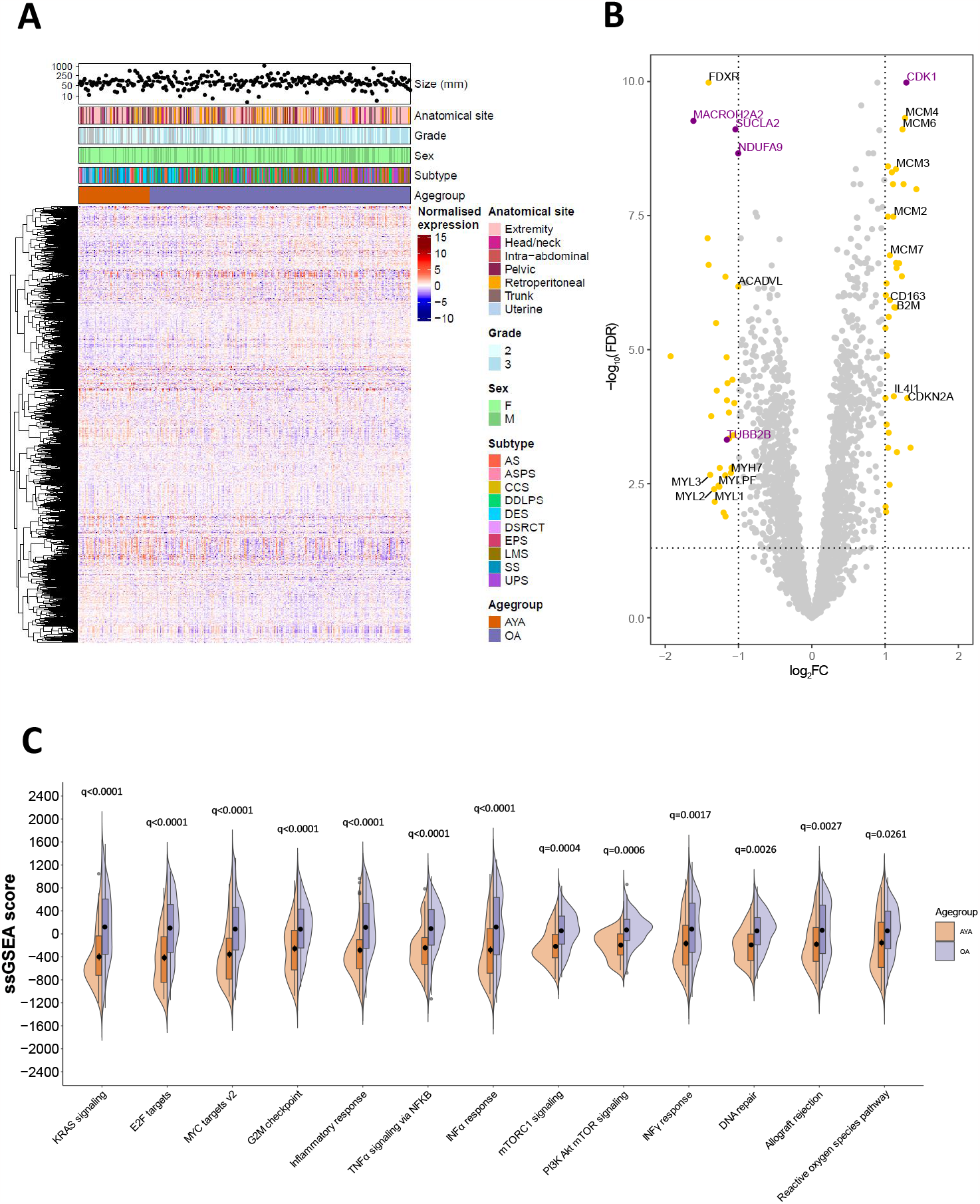
Analysis of the AYA and OA proteomic landscape. (a) Annotated heatmap showing the unsupervised clustering (Pearson’s distance) of 3299 proteins across the study cohort. Patients are ordered from youngest (left) to oldest (right). From top to bottom, panels indicate tumour size, anatomical site, tumour grade, patient sex, histological subtype, and patient age group. (b) Volcano plot showing significantly upregulated proteins in AYA and OA. Significant proteins (FDR<0.05, fold change >2) determined by multiple t-test followed by Benjamini Hochberg procedure are shown in yellow. Proteins that remained significant after multivariable logistic regression analysis are shown in purple. (c) Significantly enriched hallmark gene sets in AYA and OA patients using single sample gene set enrichment analysis (ssGSEA) scores. Significance (p<0.05) was determined by two-way analysis of variance (ANOVA) followed by Šidák correction. AS = angiosarcoma, ASPS = alveolar soft part sarcoma, CCS = clear cell sarcoma, DDLPS = dedifferentiated liposarcoma, DSRCT = desmoplastic small round cell tumour, DES = desmoid tumour, EPS = epithelioid sarcoma, LMS = leiomyosarcoma, SS = synovial sarcoma, UPS = undifferentiated pleomorphic sarcoma.

By undertaking single sample gene set enrichment analysis (ssGSEA), we show that compared to AYA patients, OA patients are significantly enriched (q<0.01) for distinct biological hallmark features (Figure 2C) including gene sets involved in cell cycle regulation (E2F targets, G2M checkpoint), oncogenic signalling (KRAS signalling, MYC targets, TNFα signalling, mTORC1 and PI3K signalling) and inflammatory pathways (inflammatory response, INFα response) (Figure 2B). AYA patients are significantly enriched for oxidative phosphorylation (q=0.004) and coagulation hallmarks (q=0.006) (Figure S2).

### Sarcoma proteomic modules highlights distinct biological pathways in the two age groups

We have previously identified 14 protein signatures based on sarcoma protein co-expression patterns termed Sarcoma Proteomic Modules (SPMs) which capture a broad spectrum of STS biology and transcend histological subtype (Table S4)^28^. We first compared the enrichment of SPMs in the AYA and OA patients and show that both groups harboured largely distinct expression levels of SPM proteins (Figure 3A). While OA patients were enriched for SPM1 (muscle system components), SPM2 and SPM4 (splicing proteins) and SPM6 (DNA replication proteins), AYA patients showed an enrichment of a much broader spectrum of biological processes. This included SPM3 (splicing proteins), SPM7 (immune proteins), SPM8 (cell adhesion & ECM proteins), SPM9 and SPM10 (vesicle transport), SPM11 (translational regulation), SPM13 (oxidative phosphorylation) and SPM14 (proteosome proteins).

**Figure 3.**
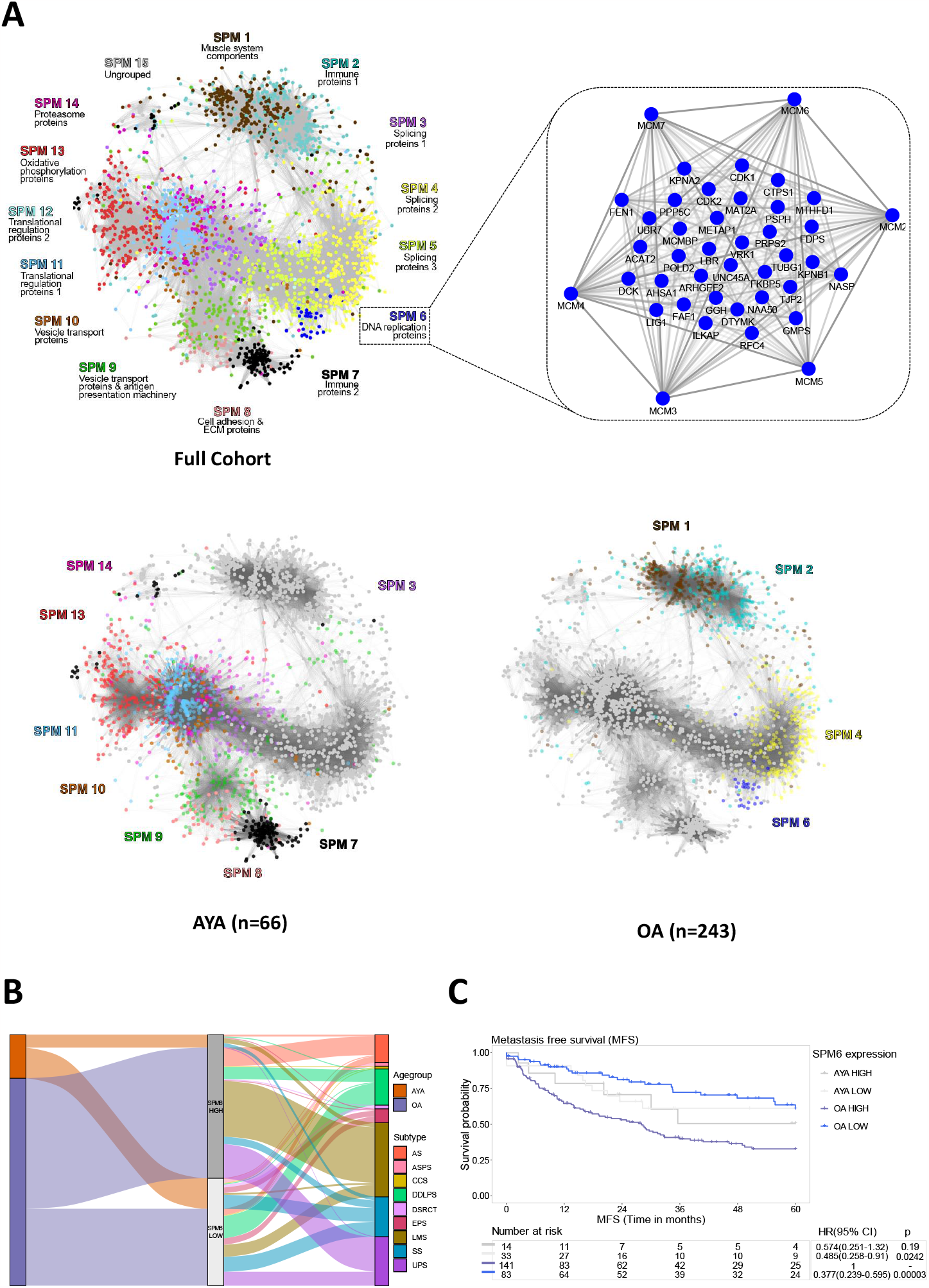
Differential expression of sarcoma proteomic modules (SPM) in AYA and OA patients. (a) Protein co-expression network showing the 14 previously described sarcoma proteomic modules (SPMs) identified in the full proteomic cohort, with a focus on the SPM6 network (Table S4). Each node represents a protein and is coloured based on SPM membership. Edges indicate correlation between protein expression, and thickness of edges are scaled to the correlation score. The protein co-expression networks for adolescent and young adults (AYA) and older adults (OA) indicate the SPMs enriched in each age group, with intensity of node colours scaled to median SPM expression level. Figure is modified from^28^ (b) Sankey plot showing the distribution of each patient age group and histological subtype that falls into the SPM6 high and low expression groups. (c) Kaplan–Meier plot of metastasis free survival (MFS) for AYA and OA patients with high and low median expression levels of SPM6 proteins. Hazard ratio (HR), 95% confidence intervals (CI) and p-value determined by univariable Cox regression. AS = angiosarcoma, ASPS = alveolar soft part sarcoma, CCS = clear cell sarcoma, DDLPS = dedifferentiated liposarcoma, DSRCT = desmoplastic small round cell tumour, EPS = epithelioid sarcoma, LMS = leiomyosarcoma, SS = synovial sarcoma, UPS = undifferentiated pleomorphic sarcoma.

SPM6 is a DNA replication module (Figure 3A) which we have shown to be prognostic for metastasis free survival (MFS) across the whole age range^28^. To further refine this candidate biomarker signature, we evaluated the histological subtype distribution of cases classified into SPM6-high and SPM6-low subgroups based on the median protein expression levels of the 41 proteins that make up SPM6 (Figure 3A and Table S4) in each of the two age groups. The Sankey plot shows that there is broad representation of histotypes in each SPM6 group, with the exception of ASPS which is only found in the SPM6-low group (Figure 3B). In addition, there was a similar distribution of AYA or OA patients in both the SPM6-high and SPM6-low groups (Figure 3B). Combining SPM6 and age stratified patients into 4 subgroups (OA-SPM6-high, OA-SPM6-low, AYA-SPM6-high and AYA-SPM6-low) (Figure 3C). While univariate Cox regression analysis found that the SPM6 module was able to stratify the OA patients into two groups with significantly different MFS outcomes (HR = 0.377, 95% CI 0.239-0.595, p=2.7X10^−5^), there was no significant difference in the AYA patients (Figure 3C). Multivariable Cox proportional hazards analysis showed that that the prognostic value of SPM6 in OA patients was independent of known prognostic factors of tumour size, grade, performance status, histological subtype and anatomical location^30,31^ (HR = 0.376, 95% CI = 0.217-0.65, p = 4.64X10^−4^) (Table S5). This analysis demonstrates that SPM6 can be used as an independent risk stratification tool to identify a subgroup of OA patients (SPM6-high) with high risk of distant relapse but has limited utility in AYA patients.

### Integrative functional genomic and proteomic analysis reveals prognostic significance of the spliceosome complex

We reasoned that biological pathways that are enriched in both CRISPR-based functional genomics and MS-based proteomics datasets may yield useful candidate drug targets and biomarkers for AYA and OA patients. Here we sought to integrate the pathway information gained from *in vitro* functional genomic data in a panel of sarcoma cell lines with the proteomic dataset generated from STT patients (Figure 4A). We first undertook an analysis of the genome-scale CRISPR-Cas9 loss-of-function screen data focusing on the NRSTS panel of cell lines within the CCLE database^32,33^. We identified 13 NRSTS cell lines where clinical information of patient age was available (AYA:n=5, OA:n=8, Table S6) and undertook gene set enrichment analysis (GSEA) using the Reactome pathway gene sets to determine the biological pathways with selective dependencies in OA compared to AYA lines (and vice versa). In parallel, we performed GSEA on the proteomic dataset of 309 STT patients to define differential pathways that are enriched in the two age groups. The top hits for gene sets significantly enriched in the OA group in both the CCLE functional genomics and patient-derived proteomic datasets were related to mRNA splicing (Figure 4B), while gene sets comprising the respiratory electron transport were significantly enriched in both datasets in the AYA patients (Figure S3).

**Figure 4.**
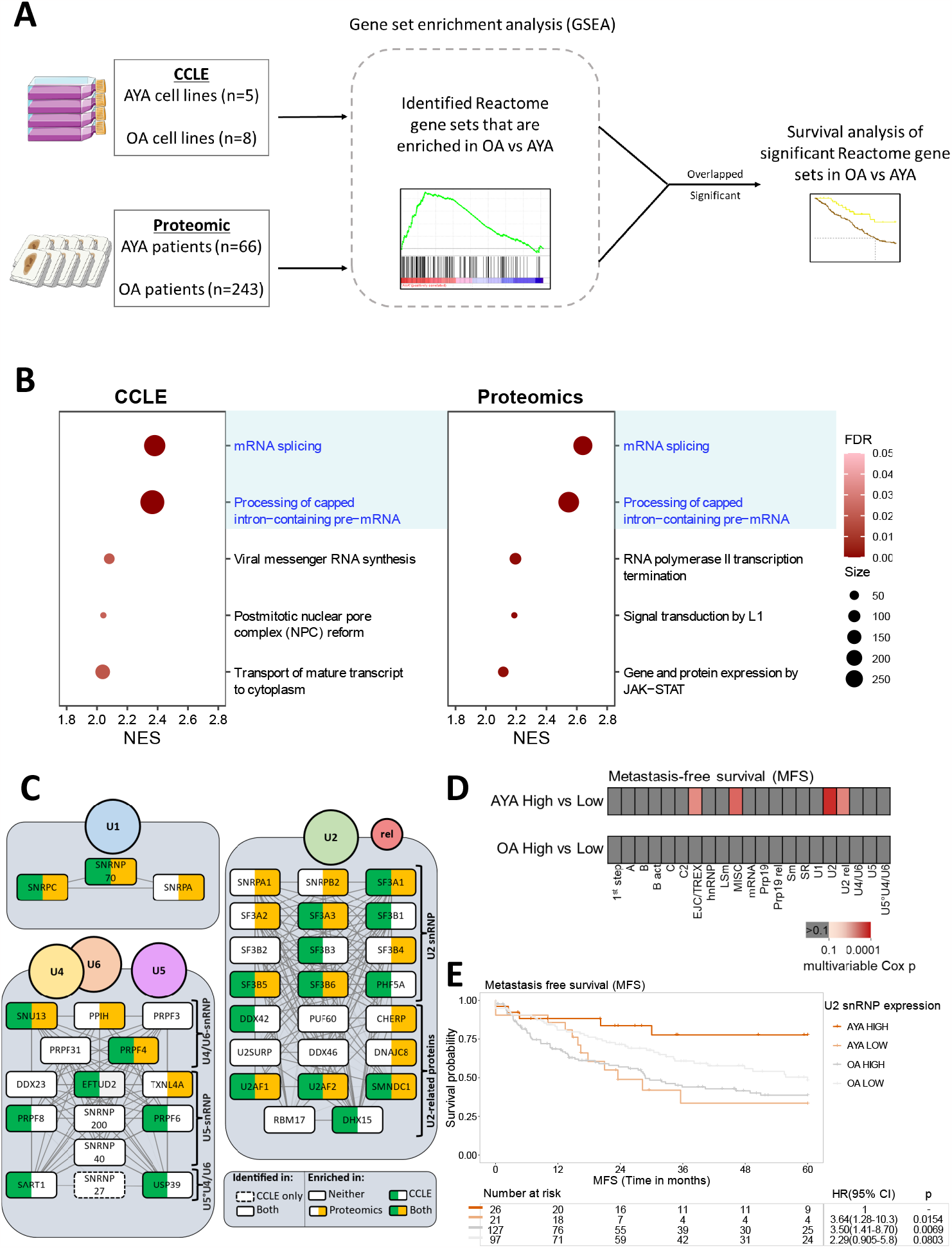
Integrative functional genomic and proteomic analysis identifies prognostic value of the spliceosome. (a) Overview schematic of the integrative functional genomic and proteomic analysis. (b) Gene set enrichment analysis (GSEA) results showing the top 5 enriched Reactome gene sets based on normalised enrichment score (NES) in the older adult (OA) group in the cell line functional genomics data from CCLE (left) and patient proteomics data (right). Overlapping gene sets enriched in both datasets are highlighted in blue. (c) Small nuclear ribonucleoprotein (snRNP) subunits of the spliceosome genes within the mRNA splicing Reactome gene set that show core enrichment in OA versus AYA patients in GSEA in the CCLE cell line functional genomic (green) and patient proteomic (yellow) dataset. Linkages indicate protein-protein interaction scores obtained from the STRING database, with a darker line indicating a higher score (cut-off >0.7). Genes that are only identified in the cell line functional genomic dataset are indicated with a dotted border. (d) Overview of multivariable Cox results for AYA and OA patients with high and low expression of each of the 21 subunits of the spliceosome complex as defined by Hegele et al (based on median protein expression), and metastasis free survival (MFS). (e) Kaplan–Meier plot of metastasis free survival (MFS) for AYA and OA patients with high and low expression of U2 snRNP proteins. Hazard ratio (HR), 95% confidence intervals (CI) and p-value determined by univariable Cox regression. FDR = false discovery rate.

Given that the OA and AYA cell lines harboured distinct dependencies on the mRNA splicing pathway genes, we hypothesized that components in this pathway may serve as prognostic signatures in the two age groups. In particular, we focused on the spliceosome complex which regulates the removal of introns from precursor mRNA during the splicing process (Figure 4C). The spliceosome is a large macromolecular complex that is comprised of >200 splicing factors that vary in their composition in a spatiotemporal manner^34^. We and others have previously shown that co-regulation of splicing factors is important in the pathology of mesenchymal and epithelial tumours^27,35,36^. As an exemplar, Figure 4C shows the proteins in the U1,U2,U4/5/6 small nuclear ribonucleoprotein (snRNP) subunits of the spliceosome complex that are enriched in OA versus AYA patients in the CCLE functional genomics as well as the proteomics datasets in the GSEA. We systematically assessed the prognostic value of proteins that make up each of the spliceosome functional component subunits as defined by Hegele et al^37^. Out of a total of 21 spliceosome subunits found in our dataset, only the U2 snRNP and a none-core miscellaneous (MISC) group of splicing factors were identified to be prognostic for MFS in AYA patients in multivariable Cox analysis (Figure 4D).

Whereas there was no significant difference in MFS between the two age groups (Figure S1), when categorised by median expression levels of U2 snRNP proteins (n=12 proteins, Figure 4C), AYA patients with high U2 snRNP expression (U2-high) had superior MFS outcomes compared to those with low U2 snRNP levels (U2-low) (multivariable: HR = 4.5, 95% CI = 1.27-15.9, p = 0.020) (Figure 4E and Table S7). No spliceosome subunit protein signatures were identified to be prognostic for OA in multivariable Cox analysis. These findings highlight the utility of integrating functional genomics data with proteomic profiling as a means of defining new prognostic factors in AYA patients.

## Discussion

AYA patients with STT are an understudied age group with disparities in treatment and survival outcomes, where improvements in 5-year survival rates over the past two decades have lagged behind other age groups^4^. Furthermore, the use of intensive multi-modal therapy often leads to chronic health conditions and secondary malignancies in AYA patients^38,39^. Rather than the current “one size fits all” approach where AYA patients, in particular those with NRSTS, are offered treatments which have been optimised in OA, tailored strategies using targeted agents and risk stratification tools could have substantial impact on survivorship and management of late effects. As a result of the under-representation of AYA patients in most molecular and biological studies in all cancer types including STT^11^, there is a poor knowledge of the biological pathways that are unique to the AYA age group which is an obstacle to developing precision medicine approaches for these patients. Notably, each of the large-scale proteomic profiling studies published thus far by The Clinical Proteomic Tumour Analysis Consortium (CPTAC) include less than 10 AYA patients^40-47^. Here we present the first large-scale analysis of the proteomic features of AYA and OA patients with STT. We show that there are inherent biological and pathway differences in the two age groups which are maintained even when confounding variables such as tumour grade, size and histological subtypes are considered. We further demonstrate that integration of *in vitro* functional genomic data in a panel of NRSTS cell lines with the patient-level proteomic data leads to the prioritisation of age-specific vulnerabilities and independent prognostic factors which provide new avenues for personalised treatment of AYA patients.

Several age-associated pan-cancer genomic analyses have shown that aging is associated with chronic inflammation and reprogramming of the immune cell landscape^48^. Our study finds that OA patients with STT are enriched in proteins involved in inflammatory response and INFα signalling hallmarks. This is consistent with a previous report by Lee et al^17^, who demonstrated using GSEA and immune cell deconvolution of transcriptomic data that sarcoma patients that are <50 years of age have lower interferon responses and lymphocyte infiltration than those >50 years. We also determined that OA patients are enriched in proteins involved in cell cycle regulation including the E2F targets and G2M checkpoint hallmarks. In agreement with our study, Chatsirisupachai et al., has shown in a pan-cancer analysis that mutations and somatic copy number alterations of genes within the cell cycle pathway are strongly enriched in tumours from older patients^16^. Interestingly, our data indicates that tumours from AYA patients harbour elevated levels of proteins involved in mitochondrial metabolism and the oxidative phosphorylation pathway which could be indicative of metabolic rewiring in younger patients. Future investigation on the functional role of metabolic rewiring in STT subtypes in this age group is warranted^49^.

A previous analysis of genome-scale CRISPR-Cas9-based loss of function screening data in a panel of paediatric cancer cell lines identified vulnerabilities that were distinct from cell lines derived from adult patients^50^, suggesting that oncology drugs that are used in adult patients may not always be applicable to childhood cancers. In this study, we focused on the pathway vulnerabilities that are specific to sarcoma cell lines in either the AYA or OA age groups and undertook analysis to compare the GSEA outputs from the genome-scale CRISPR screening data with the patient level proteomics data. Our analysis finds that *in vitro* pathway dependencies observed in sarcoma cell lines derived from patients of different age groups correspond to significantly higher expression levels of pathway proteins in either AYA (respiratory electron transport) or OA (mRNA splicing) patients. Sarcomas are a group of diseases of unmet need with a lack of effective therapies and novel agents. Investigational drugs that target components for each of these pathways are available for repurposing^51-53^ and should be evaluated in prospective studies in the different age groups. Our data further suggest that high protein expression levels of these proteins in sarcoma tissue specimens may facilitate selection of patients who are most likely to benefit from these investigational agents and therefore should be incorporated as candidate biomarkers in clinical trial design.

Despite optimal clinical management, a substantial proportion of NRSTS patients (up to 50%) with localised disease experience distant relapse following surgery^15^. Stratification of these high-risk patients has been limited to the use of nomograms which consider known prognostic factors including tumour grade, size, histological subtype and age amongst other variables^54-56^. There are currently very few molecular prognostic signatures for NRSTS and none which are optimised for AYA patients^57^. Here we show that specific subunits of the spliceosome complex are independent prognostic factors in AYA patients. In particular, AYA patients with low tumour protein expression levels of the U2 snRNP spliceosome subunit are at higher risk of developing metastasis compared to those with high expression levels. These protein signatures have potential utility as precision medicine tools to tailor more aggressive treatment strategies such as peri-operative chemo/radiotherapy in AYA patients that are predicted to have higher risk of distant relapse. Conversely, low-risk AYA patients may be spared potential overtreatment thereby reducing the risk of chronic health conditions and late effects. Mechanistically it is not clear why AYA tumours with reduced spliceosome levels appear to have more aggressive features and future functional experiments are required to dissect the role of individual splicesome protein components in AYA sarcoma cell lines. Our study further highlights the importance of performing age-specific studies to delineate biomarkers tailored for the clinical management of AYA patients.

We acknowledge several limitations of this study. This is a retrospective cohort which is prone to selection bias and therefore the study is hypothesis generating and our findings need to be validated in independent cohorts. STT comprise a broad range of histological subtypes and our study is limited to 10 and 7 histologies in the proteomic and the functional genomics datasets respectively. Future studies which include wider histological subtype representation is needed to determine if our findings are generalisable to all AYA patients, although this may be challenging given the limited number of publicly available AYA STT cell line models available for functional studies^58^. Since our study does not include a comparative analysis of proteomic profiles of normal tissue from AYA and OA individuals, we cannot exclude the possibility that some of the enriched protein signatures and pathways identified in this study are the result of physiological aging rather than being tumour specific. Despite this limitation, our data identify age-specific protein signatures with prognostic value in MFS in both AYA and OA patients which is indicative of pathological disease relevance.

In summary, we have undertaken a deep analysis of the biological differences in the proteomic profiles of STT patients in the AYA and OA age groups. We highlight important protein-specific pathways and genetic vulnerabilities that are enriched in AYA patients and identify age-specific prognostic signatures to facilitate tailored clinical management of this underserved patient group.

## Methods

### Patient cohort

The cohort is comprised of 309 patients from two centres (The Royal Marsden Hospital and National Taiwan University Hospital). Only patients that were 15 years of age or older at the time of diagnosis were included in the analysis. Retrospective collection and analysis of associated clinical data was approved as part of the Royal Marsden Hospital (RMH) PROgnoStic and PrEdiCTive ImmUnoprofiling of Sarcomas (PROSPECTUS) study (NHS Research Ethics Committee Reference 16/EE/0213) or National Taiwan University Hospital (Research Ethics Committee Reference 201912226RINB). Baseline clinicopathological characteristics and survival data were collected by retrospective review of medical records as part of our previous study^28^.

### Proteomic data

Proteomic data for this study was downloaded from ProteomeXchange (PXD036226) https://www.ebi.ac.uk/pride/archive/projects/PXD036226 ^28^. The SequestHT search engine in Proteome Discoverer 2.2 or 2.3 (Thermo Scientific, Waltham, MA, USA) was used to search the raw mass spectra against reviewed UniProt human protein entries (v2018_07 or later) for protein identification and quantification. Precursor mass tolerance was set at 20 ppm and fragment ion mass tolerance was 0.02 Da. Spectra were searched for fully tryptic peptides with maximum 2 missed cleavages. TMT6plex at N-terminus/lysine and Carbamidomethyl at cysteine were selected as fixed modifications. Dynamic modifications were oxidation of methionine and deamidation of asparagine/glutamine. Peptide confidence was estimated with the Percolator node. Peptide False Discovery Rate (FDR) was set at 0.01 and validation was based on q-value and decoy database search. The reporter ion quantifier node included an integration window tolerance of 15 ppm and integration method based on the most confident centroid peak at the MS3 level. Only unique peptides were used for quantification, considering protein groups for peptide uniqueness. Peptides with average reporter signal-to-noise > 3 were used for protein quantification. Proteins with an FDR < 0.01 and a minimum of two peptides were used for downstream analyses.

All data were processed using custom R scripts in R v4.1.1 or later. Proteins identified in <75% of samples were removed, and remaining missing values were imputed using the k-nearest neighbour (k-NN) algorithm^59^. To normalise the data and remove batch effects, data for each patient sample was divided by the corresponding reference sample and log2 transformed, followed by median centring across samples and standardising within samples. To visualise the STS proteomic dataset, hierarchical clustering was performed using Pearson correlation distance.

### CRISPR-Cas9 functional genomic data

Genome-scale CRISPR-Cas9 screening data of cell lines was downloaded from the Cancer Cell Line Encyclopaedia (CCLE) portal (https://sites.broadinstitute.org/ccle)^32^. The CRISPRGeneEffect dataset (DepMap Public 22Q4) was used for analysis. Using the model information file (https://depmap.org/portal/download/all/), only cell lines of a NRSTS subtype with a recorded patient age of >16 years were included for analysis. Cell lines derived from patients between 16-39 years old were grouped as AYA and above 39 years old as OA. Full list of included cell lines are indicated in Table S6.

### Statistical methods

All statistical tests were two-sided and where required, p values were adjusted to false discovery rate (FDR) using the Benjamini–Hochberg procedure to account for multiple comparisons (ref). Unless otherwise specified, analysis was performed using custom R scripts in R v4.1.1 or later. Two-way analysis of variance (ANOVA) and chi-square tests were implemented, with further details of statistical tests listed in the figure legends.

#### Differential expression analysis

To identify upregulated proteins in AYA and OA patients, two-tailed multiple t-test was performed and corrected for multiple comparisons by the Benjamini-Hochberg (BH) Procedure. Logistic regression analysis was performed to adjust for confounding factors of tumour size, grade, anatomical site, performance status, histological subtype, tumour margin, tumour depth. Univariate logistic regression first was performed to identify significantly different proteins between AYA and OA (FDR<0.05). Univariate logistic regression was then performed to identify significantly different confounding factors. Each significant protein’s expression was then combined with significant confounding factors and multiple logistic regression was performed with AYA and OA outcome variable.

#### Gene Set Enrichment Analysis (GSEA) and single sample GSEA (ssGSEA)

Gene Set Enrichment Analysis (GSEA) was performed using the GenePattern online tool (www.genepattern.org) to identify MSigDB Reactome gene sets (v2023.1) enriched in either age groups in the proteomic and functional genomic datasets. Single sample GSEA (ssGSEA) was similarly performed using GenePattern to score sample-specific enrichment of hallmark gene sets (v2023.1) in the proteomics dataset. ssGSEA score between AYA and OA patients were analysed using two-way analysis of variance (ANOVA) followed by Šidák correction.

#### SPM analysis and protein-protein interaction (PPI) networks

PPI networks were built in Cytoscape v3.9.11 or later. Previously described SPMs were utilised to identify proteomic signatures enriched in AYA and OA patients^28^ (Table S4). The full SPM network and individual network for SPM6 was visualised using protein co-occurrence scores and an edge weighted spring-embedded layout. For the full SPM network, a co-occurrence score threshold of >0.05 was applied. SPM expression of AYA and OA patients was calculated using the averaged median protein expression level of each SPM. The nodes in SPMs with averaged median expression <0 are shown in grey. The remaining are shown in their original colours, with the colour intensities scaled to the expression levels. To inspect the network of individual spliceosome components, protein networks were constructed using the STRING scores obtained from the STRING database v11.0, with a confidence cut off score of 0.7 and a grid layout used.

#### Survival analyses

Patients were split into -high or -low expressing groups for SPM or spliceosome components based on the median protein expression level. The association of patient groups with survival outcome were evaluated based on Kaplan Meier survival estimates and univariable Cox analysis with two-sided Wald test. Multivariable Cox analysis was used to adjust for clinicopathological variables. Three survival outcome endpoints (events) were used. Overall survival (OS) is defined as time from primary disease surgery to death from any cause. Metastasis free survival (MFS) is defined as time from primary disease surgery to radiologically confirmed metastatic disease or death. Local recurrence free survival is defined as time from primary disease surgery to radiologically confirmed local recurrence or death (local recurrence free survival/LRFS). Patients who did not have an event were censored at their last follow-up time, up to 60 months.

## Acknowledgements

This study is funded by grants from the Sarah Burkeman Trust, Cancer Research UK (C56167/A29363), Children’s Cancer and Leukaemia Group and Little Princess Trust (CCLG 2023 09), Royal Marsden Cancer Charity, The Institute of Cancer Research, and the National Institute for Health Research (NIHR) Biomedical Research Centre at The Royal Marsden NHS Foundation Trust and The Institute of Cancer Research, and a charitable donation from Geoff Crocker and Bristol Care Homes to P.H.H; Sarcoma UK (SUK03.2019) to R.L.J; Ministry of Technology and Science of Taiwan grant (109-BOT-I-002-502) to T.W.C.

## Author Contributions

Conceptualization: P.H.H.; Formal Analysis: Y.B.T, K.L., H.P, A.S.; Validation: M.C.; Data Curation: J.B., C.P.W., A.A.; Resources: T.W.C., K.T., R.L.J.; Writing – Original Draft: Y.B.T., P.H.H.; Writing – Review & Editing: All authors.; Supervision: A.S., P.H.H.; and Funding Acquisition: T.W.C, R.L.J and P.H.H.

