## Supplementary figures and images for "Proteomic Features of Adolescents and Young Adults with Soft Tissue Tumours"

### Figure S1

**A**

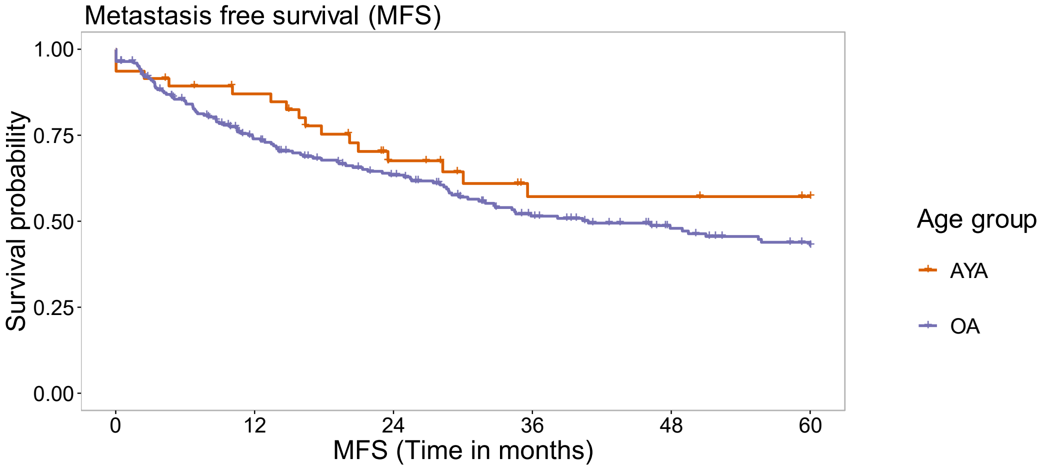


| Number at risk | | | | | | | HR(95% CI) | p |
|---|---|---|---|---|---|---|---|---|
| AYA | 47 | 38 | 23 | 15 | 15 | 13 | 0.703(0.421-1.17) | 0.178 |
| OA | 224 | 147 | 114 | 81 | 61 | 49 | 1 | - |

**B**

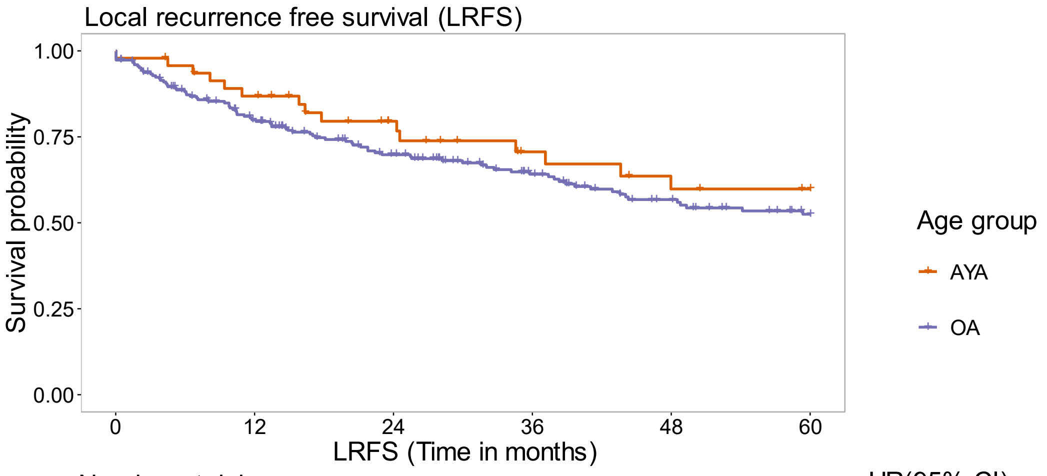


| Number at risk | | | | | | | HR(95% CI) | p |
|---|---|---|---|---|---|---|---|---|
| AYA | 47 | 39 | 28 | 20 | 16 | 14 | 0.759(0.438-1.31) | 0.324 |
| OA | 224 | 161 | 124 | 93 | 71 | 55 | 1 | - |

### Figure S2

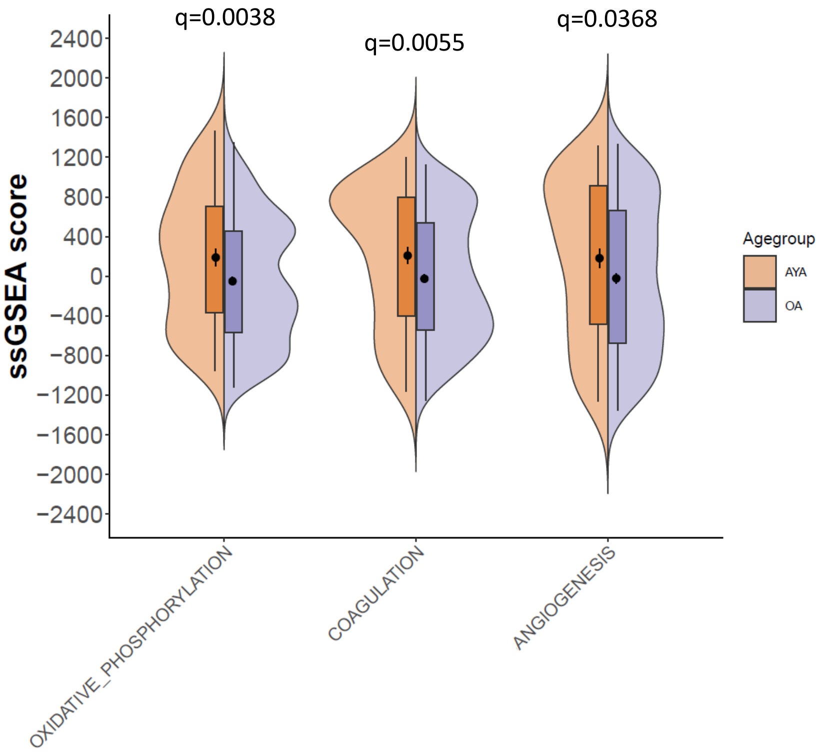

### Figure S3

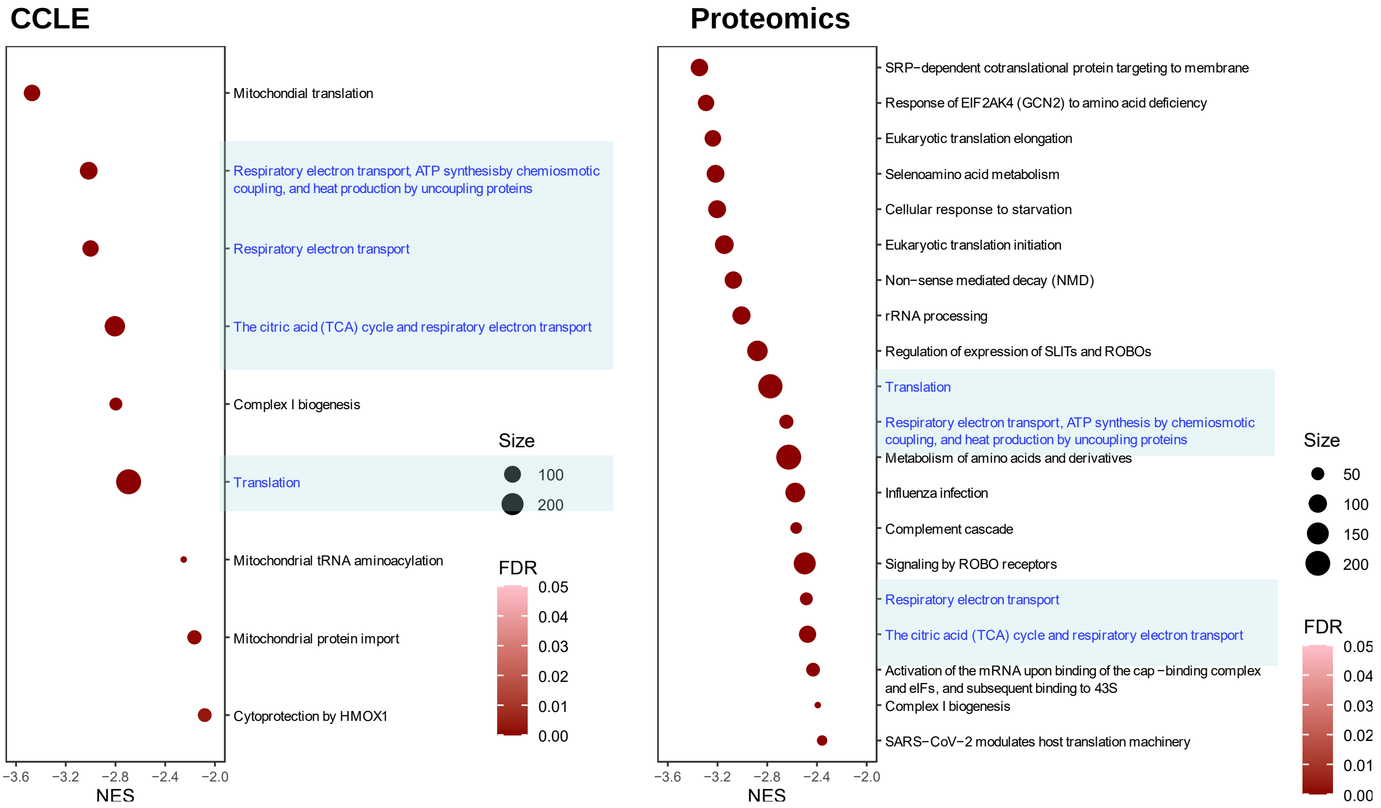
