## Supplementary material for "Proteomic Features of Adolescents and Young Adults with Soft Tissue Tumours": Table S1

**Table S1.** Clinicopathological features of the study cohort and associations with adolescent and young adult (AYA) and older adult (OA) age groups.

| **Variable** | | **Total** | **AYA** | **OA** | **Test results** | | | |
| --- | --- | --- | --- | --- | --- | --- | --- | --- |
|  |  | **n=309** | **n=66** | **n=243** | **Test performed** | **χ-squared** | **degrees of freedom** | **p value** |
| **Anatomical site [n(%)]** | **Extremity** | 124 (40.1) | 26 (39.4) | 98 (40.3) | Chi-square | 15.77 | 6 | **0.0151** |
|  | **Head/neck** | 11 (3.6) | 3 (4.5) | 8 (3.3) |  |  |  |  |
|  | **Intra-abdominal** | 25 (8.1) | 3 (4.5) | 22 (9.1) |  |  |  |  |
|  | **Pelvic** | 22 (7.1) | 9 (13.6) | 13 (5.3) |  |  |  |  |
|  | **Retroperitoneal** | 55 (17.8) | 4 (6.1) | 51 (21.0) |  |  |  |  |
|  | **Trunk** | 63 (20.4) | 18 (27.3) | 45 (18.5) |  |  |  |  |
|  | **Uterine** | 9 (2.9) | 3 (4.5) | 6 (2.5) |  |  |  |  |
| **Subtype [n(%)]** | **AS** | 30 (9.7) | 3 (4.5) | 27 (11.1) | Chi-square | 87.35 | 9 | **<0.0001** |
|  | **ASPS** | 4 (1.3) | 4 (6.1) | - |  |  |  |  |
|  | **CCS** | 3 (1.0) | 1 (1.5) | 2 (0.8) |  |  |  |  |
|  | **DDLPS** | 39 (12.6) | 2 (3.0) | 37 (15.2) |  |  |  |  |
|  | **DES** | 37 (12.0) | 18 (27.3) | 19 (7.8) |  |  |  |  |
|  | **DSRCT** | 4 (1.3) | 3 (4.5) | 1 (0.4) |  |  |  |  |
|  | **EPS** | 16 (5.2) | 8 (12.1) | 8 (3.3) |  |  |  |  |
|  | **LMS** | 80 (25.9) | 7 (10.6) | 73 (30.0) |  |  |  |  |
|  | **SS** | 43 (13.9) | 19 (28.8) | 24 (9.9) |  |  |  |  |
|  | **UPS** | 53 (17.2) | 1 (1.5) | 52 (21.4) |  |  |  |  |
| **Sex [n(%)]** | **Female** | 194 (62.8) | 49 (74.2) | 145 (59.7) | Chi-square | 4.717 | 1 | **0.0299** |
|  | **Male** | 115 (37.2) | 17 (25.8) | 98 (40.3) |  |  |  |  |
| **Tumour depth [n(%)]** | **Deep** | 250 (80.9) | 61 (92.4) | 189 (77.8) | Chi-square | 7.441 | 2 | **0.0242** |
|  | **Superficial** | 54 (17.5) | 5 (7.6) | 49 (20.2) |  |  |  |  |
|  | **unknown** | 5 (1.6) | - | 5 (2.1) |  |  |  |  |
| **Grade [n(%)]** | **2** | 115 (37.2) | 28 (42.4) | 87 (35.8) | Chi-square | 45.81 | 2 | **<0.0001** |
|  | **3** | 139 (45.0) | 10 (15.2) | 129 (53.1) |  |  |  |  |
|  | **unknown** | 55 (17.8) | 28 (42.4) | 27 (11.1) |  |  |  |  |
| **Performance status [n(%)]** | **0** | 158 (51.1) | 49 (74.2) | 109 (44.9) | Chi-square | 19.79 | 4 | **0.0005** |
|  | **1** | 82 (26.5) | 11 (16.7) | 71 (29.2) |  |  |  |  |
|  | **2** | 16 (5.2) | - | 16 (6.6) |  |  |  |  |
|  | **3** | 5 (1.6) | - | 5 (2.1) |  |  |  |  |
|  | **unknown** | 48 (15.5) | 6 (9.1) | 42 (17.3) |  |  |  |  |
| **Tumour margin [n(%)]** | **R0** | 135 (43.7) | 25 (37.9) | 110 (45.3) | Chi-square | 1.982 | 3 | 0.5762 |
|  | **R1** | 151 (48.9) | 34 (51.5) | 117 (48.1) |  |  |  |  |
|  | **R2** | 4 (1.3) | 1 (1.5) | 3 (1.2) |  |  |  |  |
|  | **unknown** | 19 (6.1) | 6 (9.1) | 13 (5.3) |  |  |  |  |
| **Log tumour size (mm) [n(%)]** | **<4** | 73 (23.6) | 18 (27.3) | 55 (22.6) | Chi-square | 12.27 | 3 | **0.0065** |
|  | **4-5** | 164 (53.1) | 38 (57.6) | 126 (51.9) |  |  |  |  |
|  | **>5** | 70 (22.7) | 8 (12.1) | 62 (25.5) |  |  |  |  |
|  | **unknown** | 2 (0.6) | 2 (3.0) | - |  |  |  |  |
