## Supplementary material for "Proteomic Features of Adolescents and Young Adults with Soft Tissue Tumours": Table S5

**Table S5.** Multivariable cox regression with two-sided Wald test assessing metastasis free survival (MFS) of adolescent and young adult (AYA) and older adult (OA) patients categorised as high and low expression of sarcoma proteomic module 6 (SPM6). HR = hazard ratio; CI = confidence interval.

|  | | | **Multivariable analysis (MFS)** | |
| --- | --- | --- | --- | --- |
| **Variable** | | **n** | **HR (95% CI)** | **p.value** |
| **SPM 6 group** | **OA HIGH (reference)** | 141 | - | - |
|  | **OA LOW** | 83 | **0.376 (0.217-0.65)** | **4.64E-04** |
|  | **AYA HIGH** | 14 | 0.697 (0.253-1.92) | 0.486 |
|  | **AYA LOW** | 33 | **0.297 (0.11-0.803)** | **0.0167** |
| **Subtype** | **LMS (reference)** | 80 | - | - |
|  | **UPS** | 53 | 1.09 (0.602-1.97) | 0.777 |
|  | **SS** | 43 | 1.09 (0.513-2.33) | 0.818 |
|  | **DDLPS** | 39 | 0.613 (0.258-1.45) | 0.267 |
|  | **AS** | 30 | **2.98 (1.23-7.23)** | **0.0156** |
|  | **EPS** | 15 | **6.75 (2.66-17.1)** | **5.92E-05** |
|  | **Other** | 11 | **4.47 (1.33-15)** | **0.0155** |
| **Anatomical site** | **Extremity (reference)** | 115 | - | - |
|  | **Pelvic** | 20 | 1.19 (0.554-2.58) | 0.65 |
|  | **Trunk** | 41 | 0.679 (0.312-1.48) | 0.328 |
|  | **Intra-abdominal** | 21 | 1.52 (0.77-3.01) | 0.227 |
|  | **Retroperitoneal** | 55 | 0.771 (0.387-1.53) | 0.458 |
|  | **Head/neck** | 10 | 1.08 (0.314-3.7) | 0.905 |
|  | **Uterine** | 9 | 1.84 (0.552-6.15) | 0.32 |
| **Sex** | **F (reference)** | 165 | - | - |
|  | **M** | 106 | 1.14 (0.745-1.76) | 0.539 |
| **Log[tumour size] (mm)** | **group4-5 (reference)** | 139 | - | - |
|  | **<4** | 65 | **0.33 (0.183-0.596)** | **2.40E-04** |
|  | **>5** | 65 | 1.08 (0.632-1.85) | 0.773 |
|  | **missing** | 2 | 0.967 (0.099-9.45) | 0.977 |
| **Performance status** | **ps0 (reference)** | 130 | - | - |
|  | **ps1** | 78 | **1.57 (1-2.46)** | **0.0493** |
|  | **ps2-3** | 21 | 1.36 (0.617-2.98) | 0.449 |
|  | **missing** | 42 | 1.55 (0.891-2.7) | 0.12 |
| **Grade** | **grade3 (reference)** | 139 | - | - |
|  | **grade2** | 114 | **0.641 (0.412-0.998)** | **0.0491** |
|  | **missing** | 18 | 0.61 (0.217-1.72) | 0.35 |
