## Supplementary material for "Proteomic Features of Adolescents and Young Adults with Soft Tissue Tumours": Table S7

**Table S7.** Multivariable cox regression with two-sided Wald test assessing metastasis free survival (MFS) of adolescent and young adult (AYA) and older adult (OA) patients categorised as expressing high and low levels of U2 small nuclear ribonucleoprotein (snRNP). HR = hazard ratio; CI = confidence interval.

**Table S7a**

|  | | | **Multivariable analysis (MFS)** | |
| --- | --- | --- | --- | --- |
| **Variable** | | **n** | **HR (95% CI)** | **p.value** |
| **U2 snRNP group** | **AYA HIGH (reference)** | 26 | - | - |
|  | **AYA LOW** | 21 | **4.5 (1.27-15.9)** | **0.0196** |
|  | **OA HIGH** | 127 | **3.62 (1.16-11.4)** | **0.0273** |
|  | **OA LOW** | 97 | 2.99 (0.903-9.89) | 0.073 |
| **Subtype** | **LMS (reference)** | 80 | - | - |
|  | **UPS** | 53 | 0.969 (0.536-1.75) | 0.917 |
|  | **SS** | 43 | 0.795 (0.384-1.65) | 0.538 |
|  | **DDLPS** | 39 | **0.321 (0.144-0.718)** | **0.00565** |
|  | **AS** | 30 | **2.7 (1.14-6.41)** | **0.0244** |
|  | **EPS** | 15 | **3.11 (1.28-7.55)** | **0.012** |
|  | **Other** | 11 | 1.94 (0.633-5.92) | 0.247 |
| **Anatomical site** | **Extremity (reference)** | 115 | - | - |
|  | **Pelvic** | 20 | 1.05 (0.48-2.3) | 0.901 |
|  | **Trunk** | 41 | 0.752 (0.351-1.61) | 0.463 |
|  | **Intra-abdominal** | 21 | 1.92 (0.979-3.75) | 0.0576 |
|  | **Retroperitoneal** | 55 | 1.1 (0.54-2.23) | 0.798 |
|  | **Head/neck** | 10 | 1.05 (0.313-3.53) | 0.936 |
|  | **Uterine** | 9 | 2.4 (0.722-7.95) | 0.154 |
| **Sex** | **F (reference)** | 165 | - | - |
|  | **M** | 106 | 1.19 (0.776-1.82) | 0.427 |
| **Log[tumour size] (mm)** | **group4-5 (reference)** | 139 | - | - |
|  | **<4** | 65 | **0.359 (0.198-0.651)** | **7.58E-04** |
|  | **>5** | 65 | 1.17 (0.685-2.01) | 0.561 |
|  | **missing** | 2 | 2.09 (0.221-19.7) | 0.521 |
| **Performance status** | **ps0 (reference)** | 130 | - | - |
|  | **ps1** | 78 | 1.41 (0.898-2.2) | 0.136 |
|  | **ps2-3** | 21 | 1.02 (0.458-2.29) | 0.955 |
|  | **missing** | 42 | 1.31 (0.765-2.26) | 0.323 |
| **Grade** | **grade3 (reference)** | 139 | - | - |
|  | **grade2** | 114 | **0.511 (0.331-0.789)** | **0.00243** |
|  | **missing** | 18 | **0.343 (0.123-0.959)** | **0.0413** |
